# Explicit representation of germline and non-germline residues improves antibody language modeling

**DOI:** 10.64898/2026.05.06.723387

**Authors:** Jeonghyeon Kim, Nathaniel Blalock, Ameya Kulkarni, Kensuke Nakamura, Philip A. Romero

## Abstract

Antibodies originate from germline templates and are diversified by somatic hypermutation, producing sequences in which conserved germline residues scaffold structure while rare non-germline (NGL) substitutions refine antigen binding. Current antibody language models (ALMs) treat all residues equivalently and inherit a germline bias that systematically down-weights functionally critical NGL mutations as statistical noise. We introduce PRISM, a germline-aware ALM that explicitly represents germline and non-germline residues as distinct token types over a factorized 53-token vocabulary. PRISM achieves state-of-the-art pseudo-perplexity in hypervariable CDRs and is uniquely positively correlated with experimental binding affinity across three deep mutational scanning landscapes on which all compared ALMs anti-correlate. The dual-vocabulary further enables property-specific controllable generation previously unattainable with entangled ALMs. NGL-directed sampling improves physics-based binding scores while GL-directed sampling preserves stability and solubility. These results establish disentangled germline/non-germline representation as a substantive advance in antibody language modeling.

## 1. Introduction

While general protein language models (pLMs) have established a foundation for biological sequence modeling (Lin et al., 2023; Madani et al., 2023), the unique generative constraints of immunoglobulins necessitate specialized antibody language models (ALMs) (Olsen et al., 2022; Shuai et al., 2023). However, even these specialized architectures face a fundamental statistical challenge: antibody generation is a hierarchical process combining static germline templates with dynamic diversification mechanisms (Murugan et al., 2012). While framework regions largely maintain conservation, functional specificity is driven by non-germline (NGL) residues from somatic hypermutation (SHM) or junctional diversity. These mutations optimize binding energy, but standard ALMs often fail to model them explicitly.

A critical bottleneck is the severe data imbalance in natural repertoires. Standard ALMs suffer from a “germline bias” since over 90% of residues remain identical to their germline progenitors, conflating evolutionary conservation with biochemical identity (Olsen et al., 2024). Consequently, standard ALMs systematically underweight NGL residues, treating critical binding-determining mutations as noise. This results in generated sequences that are structurally plausible but functionally conservative, lacking the targeted diversity required for high-affinity binding (Hie et al., 2024).

To address this, we introduce PRISM (**P**artitioning **R**esidue **I**dentity in **S**omatic **M**aturation), a framework designed to intentionally decouple germline conservation from functional variation (Figure 1A). Unlike approaches that simply fine-tune existing architectures, PRISM formalizes sequence generation as a factorized inference problem: *P*(token) = *P*(origin) × *P*(aa | origin). By utilizing a custom 53-token vocabulary that strictly separates NGL residues from germline templates, PRISM enables explicit control over the generative source. This holistic design achieves state-of-the-art pseudo-perplexity on hypervariable regions and enables controllable generation for targeted antibody design.

**Figure 1.**
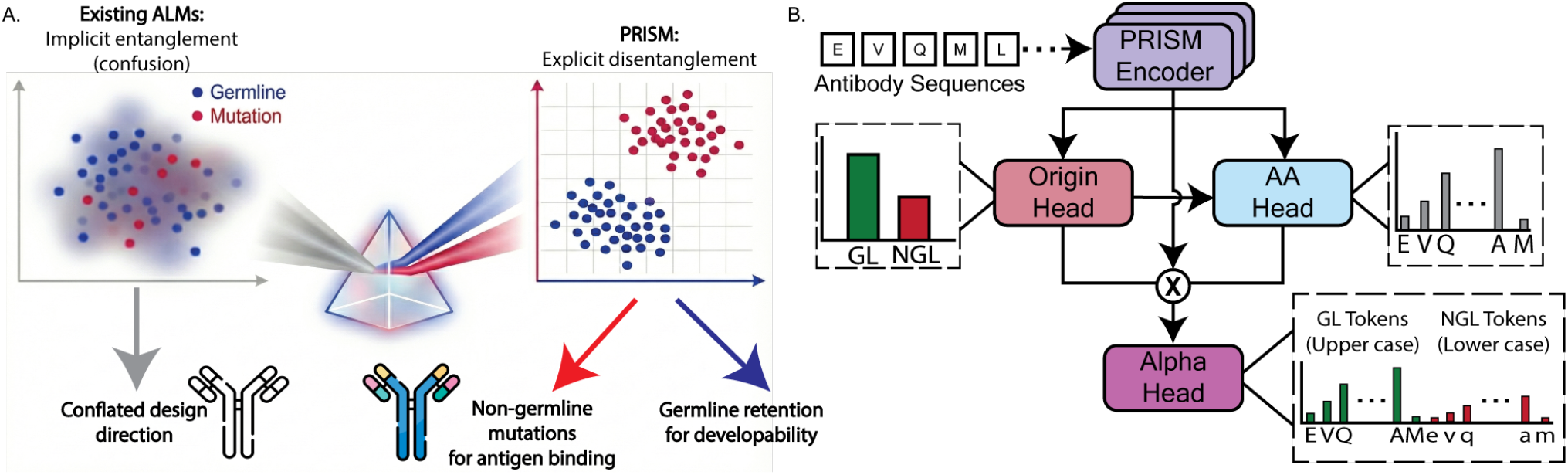
The PRISM Framework. **(A)** From Implicit Entaglement to Decomposition: Unlike existing ALMs that entangle conservation and variation, PRISM explicitly disentangles these factors. **(B)** Origin-Conditioned LM Head: PRISM employs an Origin-Conditioned LM Head in which the Origin Head first predicts the probability of NGL deviation, gating the AA Head’s predictions over a factorized 53-token vocabulary.

## 2. Preliminaries: Antibody Generation

Antibody variable domains are generated through two evolutionary phases. **V(D)J recombination** (Tonegawa, 1983) combinatorially joins V/D/J germline DNA segments to form the framework and initial CDRs against largely static templates (Chothia & Lesk, 1987). Subsequent **somatic hypermutation** (SHM) during affinity maturation (Victora & Nussenzweig, 2012) introduces point mutations, termed **non-germline (NGL)** variants, at a rate 10^6^× background (Muramatsu et al., 2000), concentrated in the CDR loops and evolutionarily selected for higher binding affinity.

### Problem formulation

Standard pLMs treat the observed sequence *x* = (*x*_1_,. .., *x*_*L*_) as a flat string, but biologically each token *x*_*i*_ is a realization of either the germline template *G* or a non-germline variant *M*:

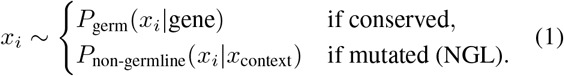

Existing models conflate these distributions. PRISM disentangles them explicitly.

## 3. PRISM Framework

Standard protein language models implicitly entangle the evolutionary origin of a residue with its physicochemical identity, leading to a strong bias toward germline sequences. Recent approaches, such as AbLang2 (Olsen et al., 2024), have attempted to mitigate this by employing reweighted loss functions (e.g., Focal Loss) to emphasize rare non-germline variants. However, these methods operate primarily at the optimization level, leaving the underlying representation entangled.

PRISM advances this paradigm by enforcing a fundamental disentanglement at the *representational* level. By structurally separating evolutionary origin from biochemical identity, PRISM enables emergent controllability over the generation process. This allows us to treat evolutionary divergence not merely as a prediction target, but as a tunable constraint. We adopt the architectural specifications of ESM-2 (Lin et al., 2023) (35M parameters, 12 layers) as our backbone, replacing its standard masked-LM head with our novel **Origin-Conditioned LM Head** — a three-component output module (Origin Head, AA Head, Alpha Head) in which the Origin Head conditions the AA Head and an Alpha Head learns to gate the evolutionary signal (Figure 1B).

### 3.1 Factorized Vocabulary

Based on the high-confidence non-germline mutation signals curated in our dataset (Appendix A), we construct an extended vocabulary 𝒱_ext_ of size 53 to support disentanglement (Appendix B.2). Uppercase tokens represent germline (GL) residues matching the template, while lowercase tokens denote non-germline (NGL) variants (2). This distinction allows the model to structurally separate evolutionary origin from biochemical identity.

### 3.2 Origin-Conditioned LM Head

Let **H** ∈ ℝ^*L*×*d*^ denote the output of the encoder, where *L* is the sequence length and *d* = 480 is the embedding dimension. The Origin-Conditioned LM Head is a three-component output module — Origin Head, AA Head, Alpha Head — in which the evolutionary prediction explicitly conditions the biochemical prediction.

#### Origin Head (Evolutionary Prior)

First, we predict the probability of non-germline status at position *i*. The Origin Head projects the contextual embedding **h**_*i*_ to a scalar logit *o*_*i*_:

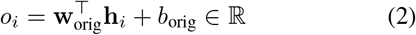

The probability of a non-germline (NGL) origin is given by π_*i*_ = *σ* (*o*_*i*_), where σ refers to sigmoid function.

#### Amino Acid (AA) Head

This head predicts the fundamental biochemical identity of the residue given the evolutionary context defined by 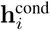. It projects the embedding into the base vocabulary space:

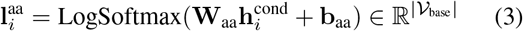

#### Gradient-Detached Conditioning

To ensure the Origin Head is optimized solely by evolutionary signals, we apply a stop-gradient operator (SG) before conditioning the downstream heads:

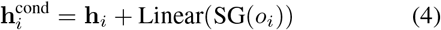

Here, SG(·) acts as an identity function during the forward pass but blocks error propagation during the backward pass. We empirically verify in Appendix K that this prevents the AA Head from delegating NGL prediction to the Origin Head, which results in a ∼2× improvement in validation NGL perplexity and does not introduce early-training instability relative to ablation experiments.

#### Alpha Head (Task-Relevance Gating)

To modulate the influence of the evolutionary prior, we introduce a scalar gating mechanism. This head predicts a position-specific mixing coefficient α_*i*_, representing a position-specific gate on the evolutionary prior:

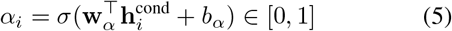

Although the Origin Head already predicts π_*i*_, a separate Alpha Head is architecturally necessary as it absorbs the gradient pressure from the language modeling loss (thereby preserving the Origin Head as a pure evolutionary classifier) and admits the evolutionary prior independently of Origin Head accuracy. Appendix J shows this is isomorphic to a position-specific tempered posterior whose tempering exponent is *learned end-to-end*, and that empirically *α*_*i*_ behaves as a *task-relevance gate* — saturating in hypermutation-prone CDRs and collapsing at IMGT-conserved structural anchors (Cys23, Trp41, Cys104, Phe/Trp118) — rather than a confidence decay on the Origin Head.

### 3.3 Alpha-Gated Logit Construction

The final distribution over the 53-token vocabulary is constructed via multiplicative gating in log-space. For any target token *y*, let ℳ (*y*) be its biochemical identity index and 𝒵 (*y*) ∈ {*GL, NGL}* be its evolutionary class. The un-normalized logit *s*_*i,y*_ is formulated as:

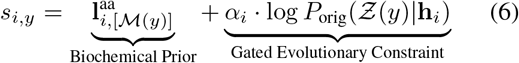

where *P*_orig_(*NGL*|**h**_*i*_)= π_*i*_ and *P*_orig_(*GL*|**h**_*i*_)= 1 - π_*i*_. This mechanism admits the Origin pathway only where it carries incremental information for residue identity, collapsing to the marginal AA distribution at structurally constrained sites, and provides a controllable knob for evolutionary exploration. Equation 6 implies a natural Bayesian interpretation as a position-specific tempered posterior, which we analyze formally in Appendix J. Finally, we apply temperature scaling with a factor *T* = 0.5 to sharpen the distribution for therapeutic engineering applications:

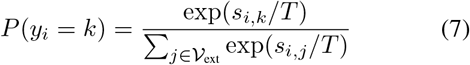

### 3.2 Training Objective and Data

#### Multi-head training objective

PRISM is trained with a weighted combination of three masked-token losses corresponding to the architectural decomposition described in Sec. 3.1–3.3: a *final-distribution loss* ℒ_final_ over the 53-token alpha-gated logits (Eq. 6), an *AA Head loss* ℒ_AA_ over the 20-canonical amino-acid prediction, and an *Origin Head loss* ℒ_origin_ over the binary GL/NGL label,

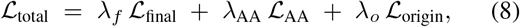

with λ_*f*_ = 2.0, λ_AA_ = 1.0, λ_o_ = 1.5. The two amino-acid losses (ℒ_final_, ℒ_AA_) are computed as a per-token-weighted focal loss (Lin et al., 2017; Olsen et al., 2024) over masked positions; to counteract the severe germline imbalance (> 90% of residues are GL), every NGL position receives a per-token weight *α*_NGL_ = 3.0 while GL positions retain weight 1.0, applied *inside* the focal-loss reduction so that gradients from rare somatic substitutions are amplified at the token level rather than at the sequence level. This per-token reweighting is the loss-side counterpart to the dual-vocabulary disentanglement: it prevents the dominant germline signal from washing out the rare NGL events whose modeling fidelity ultimately drives binding-affinity prediction (Sec. 6.1). Crucially, this loss-side reweighting is *not* what drives the representational disentanglement reported in Sec. 7: Ablation 2 retains the identical focal loss with *α*_NGL_ = 3.0 reweighting and the extended 53-token vocabulary while replacing only the Origin-Conditioned LM Head with a simple LM head, yet collapses to near-random GL/NGL separability (ARI 0.009 vs. PRISM Full’s 0.162) — the architectural factorization is separable from, and not substitutable by, the loss reweighting.

#### Region-aware masking

Standard 15% uniform masking under-samples the hypervariable loops where NGL substitutions concentrate. We therefore apply region-conditional masking probabilities – 50% on CDRs, 30% on framework regions, and 15% background – so the model is forced to reconstruct the very positions whose supervision signal is statistically rarest, again coupling the training distribution to the GL/NGL contribution PRISM is designed to recover (full schedule and ablation in Appendix C).

#### Data

We train on antibody sequences from the Observed Antibody Space (OAS) database (Kovaltsuk et al., 2018). Two corpora are used in a two-stage curriculum: an *unpaired* corpus of ∼58M heavy and ∼8.5M light chains for pretraining, and a higher-quality *paired* corpus of 619,675 heavy/light pairs for finetuning. To mitigate germline bias at the dataset level, we exclude near-germline “noise-floor” sequences using a chain-specific naive-B-cell mutation threshold (τ_HC_ = 3, τ_LC_ =2 corresponding to the 90th percentile of mutation counts in naive paired sequences), shifting the training distribution toward sequences carrying meaningful somatic variation while retaining > 80% of memory B-cell sequences (Appendix Fig. S1). Train/validation/test splits are constructed via Linclust (Steinegger & Sö ding, 2018) clustering at 100% CDR3 identity and 95% whole-sequence identity, with whole clusters assigned to a single split, eliminating clonally related leakage between train and test (test set ∼ 22,600 sequences). All hyperparameters, gene/region embedding details, and the full filtering pipeline are deferred to Appendix A–C.

#### Downstream evaluation data

We evaluate zero-shot binding affinity on three Deep Mutational Scanning (DMS) sets spanning a broad complexity spectrum: the single-mutant scan of **G6.31** (*N* = 4,274, anti-VEGF, *K*_*d*_ ≈ 0.4 nM (Koenig et al., 2017a)), the ∼16-site combinatorial **CR9114** (*N* = 65,093, anti-influenza HA (Phillips et al., 2021)), and the ∼10-site combinatorial **Trastuzumab** (*N* = 36,496, anti-HER2 (Mason et al., 2021)); plus the 41-antibody **FLAb2** binding panel (Chungyoun & Gray, 2025) for breadth. Developability is evaluated on the 246-antibody Arsiwala/Marks biophysical panel (Arsiwala et al., 2025) (Self-Interaction, Hydrophobicity, Thermal Stability, Polyreactivity, Expression, Immunogenicity), clinical Anti-Drug-Antibody response rates (Marks et al., 2021), and the FLAb2 developability assays (DSC, AC-SINS, HEK, PSR, ADA). For controllable generation (Sec. 5), the same three DMS antibodies (G6.31, CR9114, Trastuzumab) provide WT structural templates (PDB 2FJH, 4FQI, 1N8Z) for Rosetta interface and stability scoring.

## 4. Disentanglement Improves Antibody Modeling

We evaluate PRISM against general pLMs (ESM2-35M, ESM2-650M (Lin et al., 2023)) and specialized ALMs (the masked AbLang2 (Olsen et al., 2024), AntiBERTy (Ruffolo et al., 2021), Sapiens (Prihoda et al., 2022), and the autoregressive generative IgLM (Shuai et al., 2023)), and show that PRISM’s factorized architecture jointly delivers explicit GL/NGL disentanglement and superior predictive performance on functional residues.

### 4.1. Explicit Separation in Latent Space

Linear probing on frozen residue embeddings (Fig. 2A) shows PRISM achieving near-perfect GL/NGL separability (PR-AUC 0.980, F1 0.896; Fig. 2B), substantially above ESM2-35M (PR-AUC 0.354), AntiBERTy (0.929), and the generation-oriented IgLM (0.324, F1 0.312); the zero-shot Origin Head (0.958) already rivals trained probes. UMAP projections after residue-type mean centering (Appendix D) confirm this geometrically (Fig. 2C): PRISM exhibits a clear GL/NGL cleavage (ARI 0.172), whereas baselines remain entangled (ESM2-35M 0.009, IgLM 0.004).

**Figure 2.**
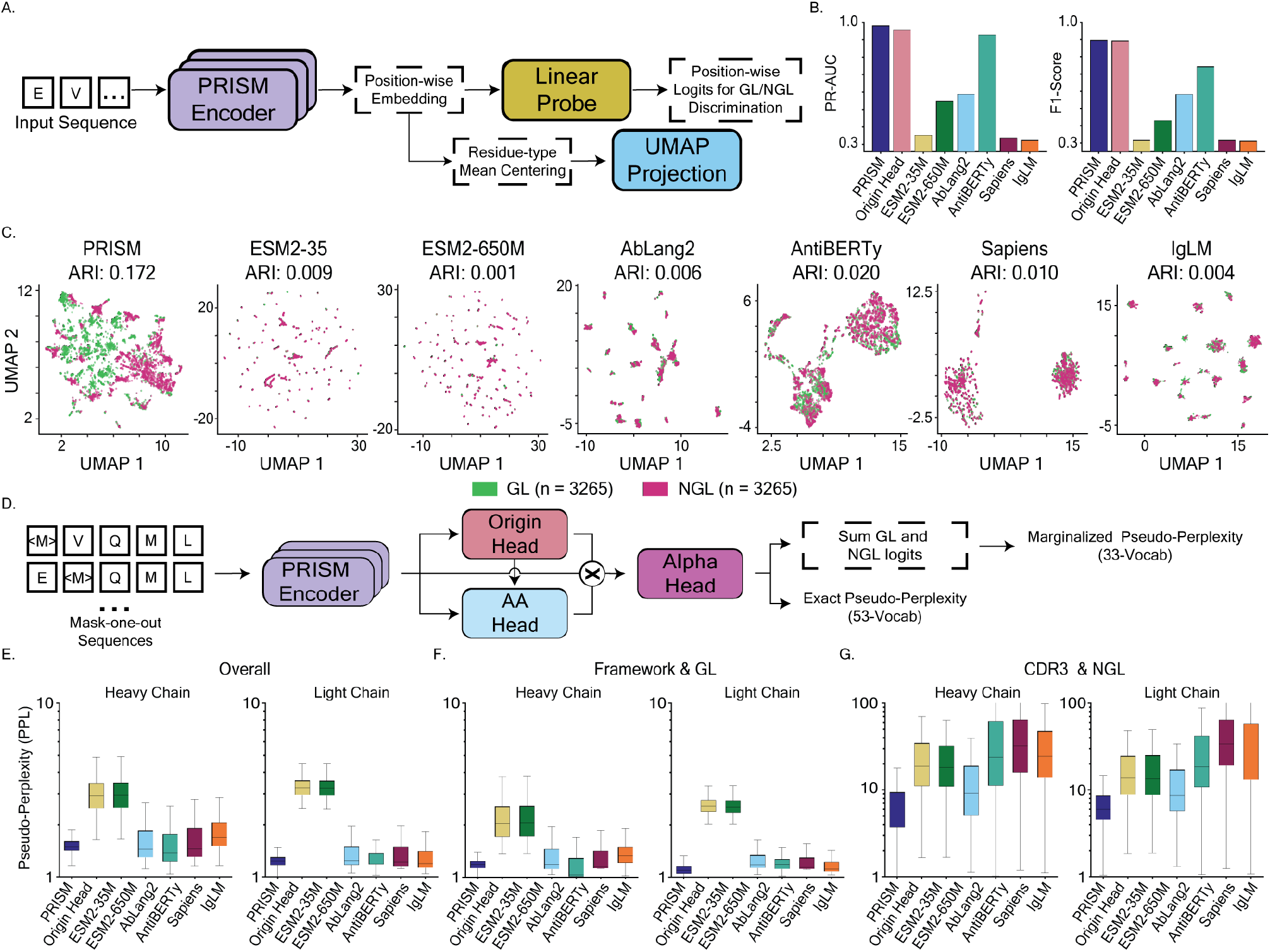
PRISM achieves explicit disentanglement and superior generative performance. **(A-C)** Linear probing and UMAP projections demonstrate that PRISM achieves near-perfect separation of GL and NGL residues (PR-AUC 0.980, ARI 0.172), whereas both masked ALMs and the generative IgLM show entangled distributions (ARI ≥ 0.020). **(D-G)** Generative evaluation via Pseudo-perplexity confirms this structural advantage. **(E)** PRISM achieves the lowest marginalized PPL overall across heavy and light chains. **(F)** While maintaining parity on stable Framework regions, **(G)** the model dramatically outperforms baselines on hypervariable CDR3-NGL residues, including the generation-specialized IgLM (6.15 vs. 25.12 on heavy chain).

### 4.2. Sequence Modeling Performance Stratified by Structure and Origin

We assess generative capability via Marginalized PPL (summing GL+NGL probabilities for fair comparison with standard-vocabulary baselines) and Exact PPL (Fig. S3E), which penalizes origin misclassification. On whole sequences (Fig. 2E) PRISM attains the lowest marginalized PPL on both heavy (1.54) and light (1.25) chains, beating ESM2-35M (∼ 3.0), AbLang2 (∼ 1.6), and IgLM (1.73/1.26). On the hypervariable CDR3-NGL residues that drive specificity, PRISM reaches 6.15 on the heavy chain versus AntiBERTy 24.08, AbLang2 9.26, and IgLM 25.12 – the last being particularly notable since IgLM is explicitly optimized for antibody generation yet still struggles on the loops defining specificity. On conserved framework regions, PRISM maintains parity with specialized baselines (∼ 1.18), confirming that disentanglement enables targeted CDR diversity without compromising scaffold developability (full breakdown in Appendix F).

## 5. Controllable Generation via Disentangled Logits

PRISM’s disentangled representation also yields a directly controllable generative interface: by selecting which slice of the alpha-gated 53-token logit drives sampling, the same model is steered toward distinct biophysical objectives without task-specific finetuning. **Protocol**. For each WT antibody, we identify the 20 positions whose WT residue receives the lowest PRISM logit – the positions most amenable to substitution – and for each generated variant, draw 10 of them without replacement, mask each, and resample from the masked-LM distribution under one of three channels: **PRISM-Full** (Eq. 6), **PRISM-NGL** (NGL tokens only), or **PRISM-GL** (GL tokens only). Baselines (ESM2-35M, AbLang2, IgLM) are generated from their native distributions under the same position selection. Variants (*n*_mut_ = 10, 100 per antibody per method) are scored along four axes: Rosetta interface ΔΔ*G* (physics-based binding), a ridge-based predictor trained on each antibody’s DMS data (sequence-based binding), Rosetta antibody-only ΔΔ*G* (stability), and CamSol pH 7.0 (solubility). The full methodology is in Appendix E.

The dual-logit decomposition surfaces a clean separation of regimes (Fig. 3B–D): **PRISM-NGL** produces the strongest binding-favorable variants on the Rosetta interface metric across all three antibodies, while **PRISM-GL** dominates on Rosetta stability and CamSol. For G6.31, where the site-saturated DMS renders the ridge-based predictor most reliable, PRISM-NGL leads on both the ridge-based predictor and Rosetta interface, providing the cleanest evidence that NGL-biased sampling captures genuine binding-relevant variation.

**Figure 3.**
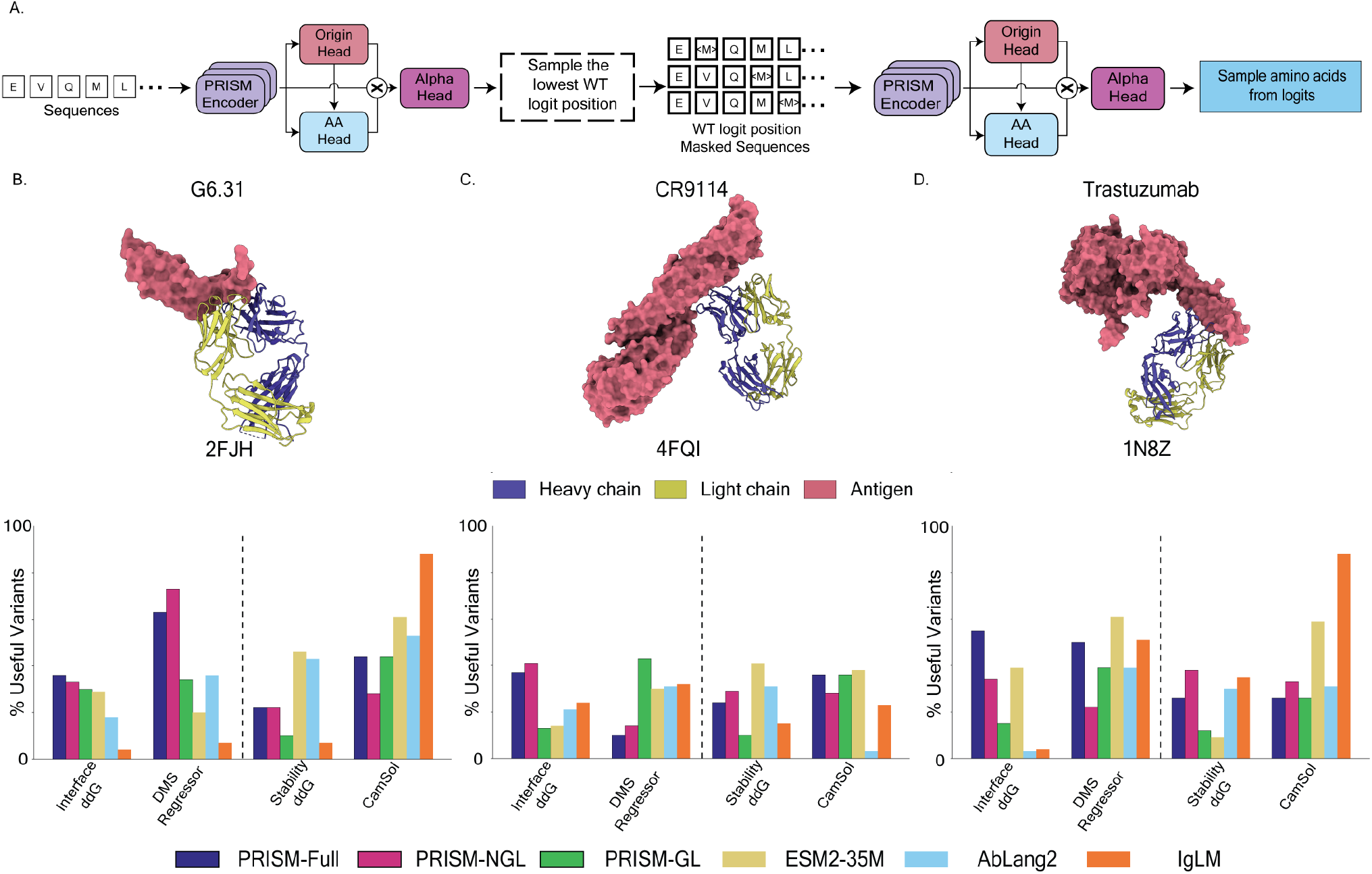
Disentangled-logit generation enables property-specific control. **(A)** Protocol: for each WT antibody we take the 20 positions with the lowest WT logit under PRISM, draw 10 per variant, mask, and re-sample under one of three channels – PRISM-Full (alpha-gated), PRISM-NGL (NGL only), PRISM-GL (GL only). **(B–D)** Fraction of generated 10-mutation variants improving over WT on each axis for G6.31 (2FJH), CR9114 (4FQI), and Trastuzumab (1N8Z). PRISM-NGL leads on Rosetta interface ΔΔ*G* (binding); PRISM-GL leads on Rosetta stability and CamSol (solubility); PRISM-Full compares favorably against all generative baselines (ESM2-35M, AbLang2, IgLM) on the aggregate. The DMS regressor axis is a DMS-trained ridge regression binding affinity predictors; G6.31 alone has site-saturated DMS coverage (Appendix E.4).

For CR9114 and Trastuzumab, the ridge-based predictor ranks PRISM-NGL slightly below some baselines, but the Rosetta interface signal, independent of the DMS coverage limitations, still places PRISM-NGL on top, and the PRISM-GL advantage on stability and CamSol is consistent across all three antibodies; per-variant distributions and spatial mutation patterns (Appendix Figs. S4, S5) confirm this geometrically, with PRISM-NGL concentrating substitutions in CDRs and PRISM-GL spreading them through framework-adjacent sites.

**PRISM-Full** lies between these specialized modes but compares favorably against all generative baselines on the aggregate of binding, stability, and solubility. Together, these results establish PRISM as a controllable generator with two practically useful modes—NGL for affinity and GL for developability—and a strong unconstrained default.

## 6. Zero-Shot Prediction of Binding Affinity and Developability

All evaluation datasets are introduced in Section 3.4. We score binding affinity via NGL probability and developability via GL probability (Jain et al., 2017), sign-adjusting all Spearman correlations (*ρ*) so positive values denote favorable outcomes.

### 6.1 Binding Affinity Prediction

PRISM is uniquely positively correlated with experimental affinity on *every* landscape tested (Fig. 4B–D). On the combinatorial **CR9114** (*ρ* = +0.391, 4.6× enrichment at top-10%), every baseline correlates *negatively* (*ρ* ∈ [−0.428, −0.326]) and yields anti-enrichment (0.1× −0.4×), making PRISM the only method delivering practical screening guidance in this regime. On **Trastuzumab**, PRISM (*ρ*= +0.327, 2.6 ×) edges out the strongest specialized ALM (AntiBERTy: +0.297, 2.5×). On the strongly affinity-matured **G6.31** (*K*_*d*_ ≈ 0.4 nM), PRISM is the only model with a positive Spearman (+0.158), while every baseline yields negative ranks (Sapiens −0.018 down to ESM2-35M −0.202); G6.31’s narrower enrichment dynamic range (2.3×, comparable to IgLM’s 2.6× despite IgLM’s *ρ* = −0.065) is the expected consequence of scoring near a local affinity optimum, and PRISM’s advantage is clearest in rank correlation (Appendix H.3). On **FLAb2** (Fig. 4E–G), PRISM has the strongest Spearman on 41.5% of antibodies (next-best ESM2-650M: 19.5%), the lowest mean rank (3.59 vs. all baselines > 4.00), and the highest median per-antibody Spearman (0.091).

**Figure 4.**
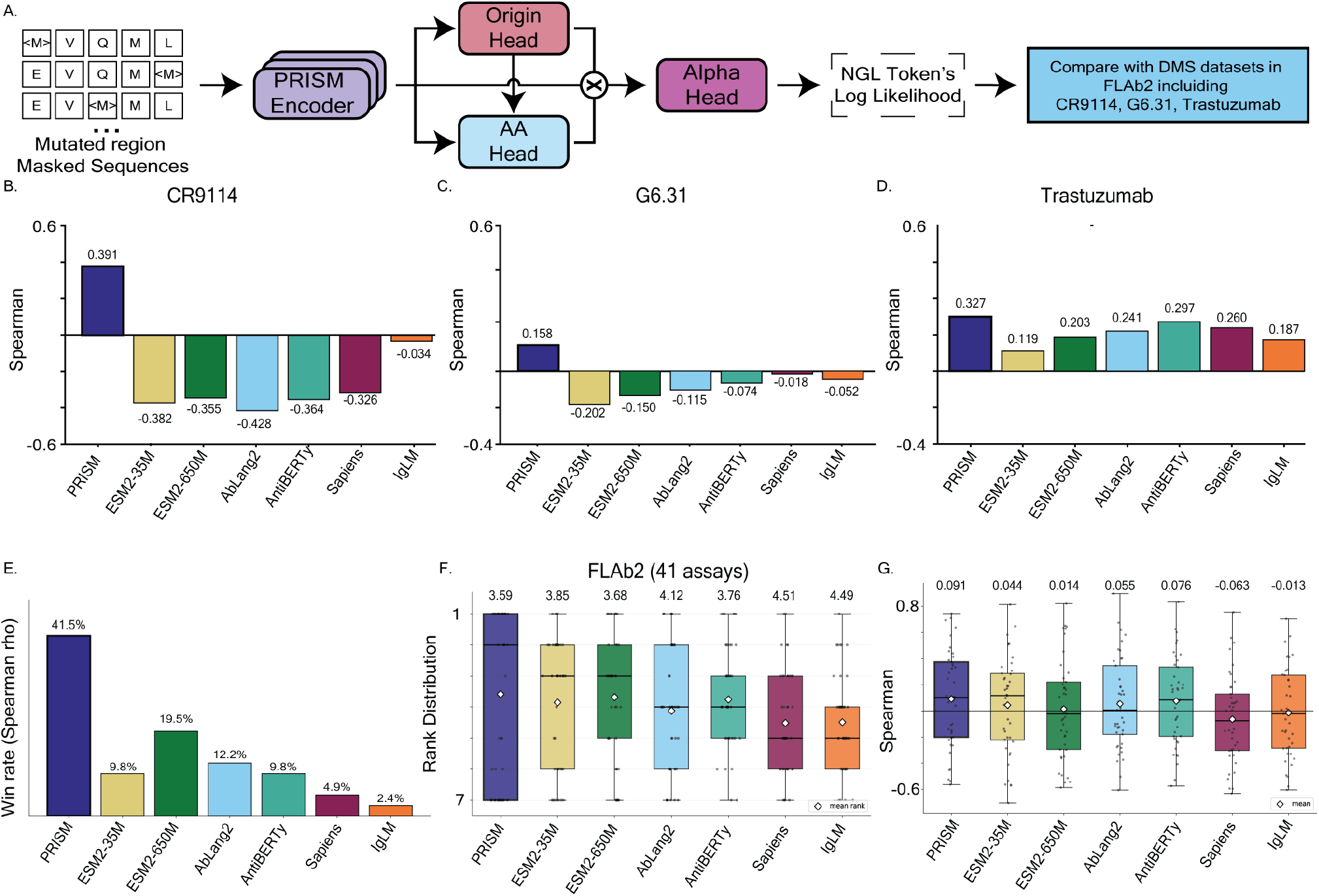
Zero-shot prediction of binding affinity. **(A)** Mask-one-out evaluation: PRISM’s NGL-constrained log-likelihood vs. experimental affinity on three DMS sets and the 41-antibody **FLAb2** panel. **(B–D)** Spearman *ρ* on CR9114 (*N* = 65,093), G6.31 (*N* = 4,274), Trastuzumab (*N* = 36,496): PRISM is the *only* method positive on all three (CR9114 +0.391, G6.31 +0.158, Trastuzumab +0.327); every baseline is negative on CR9114 ([−0.428, −0.326]) and G6.31 ([−0.018, −0.202]), indicating systematic germline bias. **(E–G) FLAb2 (41 antibodies):** PRISM has the top per-assay win rate (41.5% vs. ESM2-650M 19.5%), the lowest mean rank (3.59 vs. all baselines > 4.00), and the highest median per-antibody Spearman (0.091 vs. AntiBERTy 0.078, ESM2-650M 0.026). See Sec. 6.1, Appendix H.3.

### 6.2. Developability Assessment

PRISM delivers a balanced developability profile (Fig. 5). On the Arsiwala/Marks 6-trait panel (Appendix Fig. S6A– F), PRISM ranks **first** on Self-Interaction, Hydrophobicity, and Thermal Stability, with positive correlations across all six traits and 2nd/4th on Immunogenicity/Expression. On the broader **FLAb2** developability panel (Appendix Fig. S6G–K), PRISM leads on DSC (thermal stability) and AC-SINS (aggregation) and stays competitive on HEK, PSR, and ADA – a state-of-the-art stability profile without compromising reactivity or immunogenicity. Pearson analysis (Appendix Fig. S7) reproduces the same ordering.

**Figure 5.**
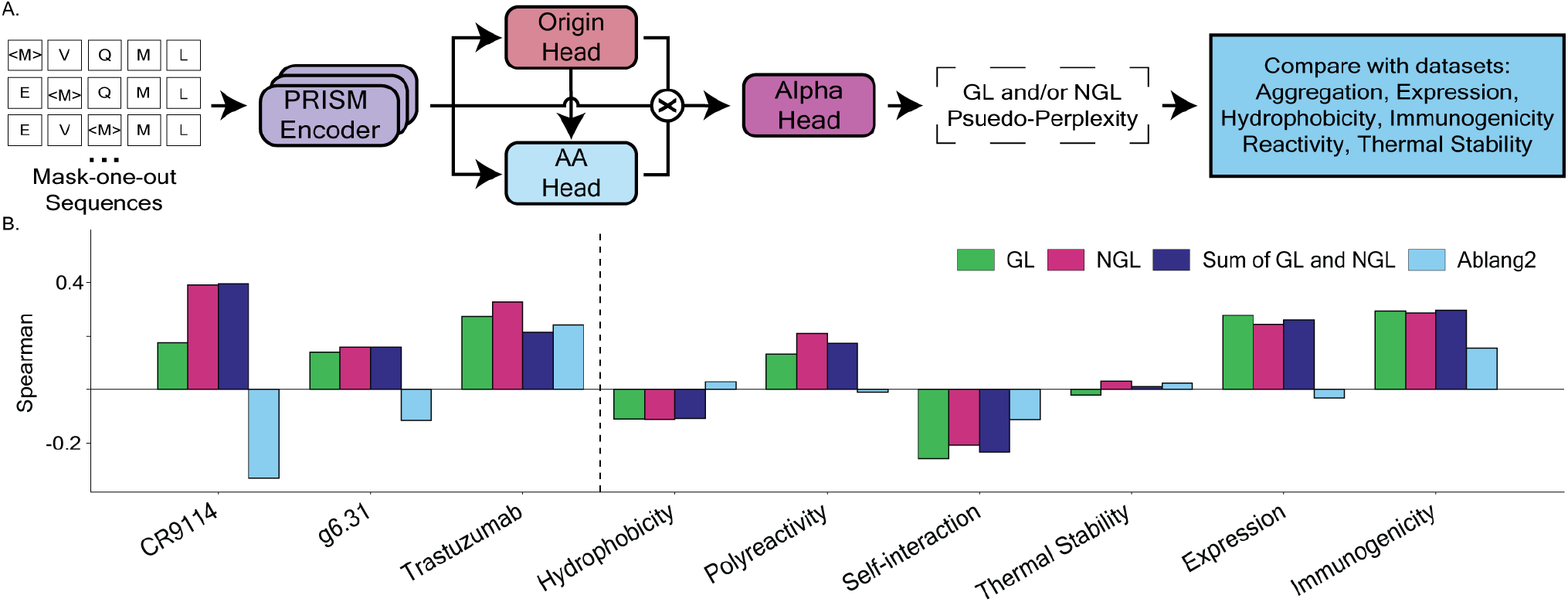
Disentanglement of Function and Developability. **(A)** Schematic: mask-one-out sequences are scored by PRISM’s GL-, NGL-, and Sum-of-(GL,NGL) log-likelihoods (via the Origin Head → AA Head → Alpha Head pipeline) and compared with binding-affinity DMS sets and biophysical assays. **(B)** Zero-shot Spearman of GL (Green), NGL (Magenta), and Marginal/Sum (Navy) scores against AbLang2 (Light Blue) across the binding-affinity DMS sets (left of dashed line: CR9114, g6.31, Trastuzumab) and six developability traits (right of dashed line). On binding affinity, GL and NGL are clearly separated, with NGL consistently outperforming GL — the gap widest on CR9114, where AbLang2 collapses to a strongly negative correlation while both PRISM scores remain positive. On developability, the GL and NGL channels are largely comparable across most traits; the exception is on Expression and Immunogenicity — the two developability traits most directly tied to germline conservation — where GL carries slightly more signal than NGL, in line with the biological role of germline identity in determining expression yield and ADA risk. The Marginal score (Navy) frequently degrades performance below the better-aligned individual score (e.g., on Trastuzumab and CR9114), consistent with a “signal cancellation” effect when entangling these complementary axes in a single distribution.

### 6.3. Validation of Dual-Vocabulary Strategy: The Necessity of Disentanglement

To test the premise that NGL mutations drive function while GL identity scaffolds developability, we decomposed PRISM’s output into **GL-only, NGL-only**, and **Marginal** (sum) scores and compared each against AbLang2 (Fig. 5B). On *binding affinity*, NGL (Magenta) consistently outperforms GL (Green) across CR9114, G6.31, and Trastuzumab, with the gap widest on CR9114 where AbLang2 (Light Blue) collapses to a strongly negative correlation while both PRISM scores remain positive – the dual-vocabulary decomposition cleanly recovers the SHM-driven affinity signal where AbLang2 cannot. On developability, GL and NGL are largely comparable across Hydrophobicity, Self-Interaction, Polyreactivity, and Thermal Stability, but on the two traits most directly tied to germline identity – **Expression** (yield is determined partly by germline V-gene) and **Immunogenicity** (framework similarity to germline shapes ADA risk) – the GL score carries slightly more information than NGL.

The **Marginal** score (Navy) – mimicking standard pLMs that do not separate origin – frequently underperforms the better-aligned specialized score (e.g., on Trastuzumab and CR9114), consistent with a “signal cancellation” effect when complementary axes are entangled in a single distribution. Together, (i) the clear GL/NGL split on binding, (ii) the biologically aligned GL advantage on Expression and Immunogenicity, and (iii) the cost of marginalizing the two channels establish PRISM as a controllable generator that can independently modulate developability and binding – a capability unattainable by entangled models.

## 7. Ablation Studies

We ablate three orthogonal axes – architecture (multi-head vs. simple head), training (pretrained vs. scratch), and initialization (random vs. ESM2) – via four variants (Fig. 6A): **Ablation 1** drops unpaired pretraining while keeping the multi-head; **Ablation 2** keeps pretraining but uses a simple LM head; **Ablation 3** drops both; and **PRISM-less** fine-tunes ESM2-35M on antibody data with a simple head over the standard (upper-case) vocabulary – the strongest “naive transfer” baseline. All variants share the identical training pipeline — focal loss with per-token NGL reweighting *α*_NGL_ = 3.0, region-aware masking, and the hyperparameters of Sec. C — and Ablations 1–3 additionally share the extended 53-token vocabulary; variants differ *only* in the axes named above, isolating the architectural and pretraining contributions from the loss-side reweighting mechanism.

**Figure 6.**
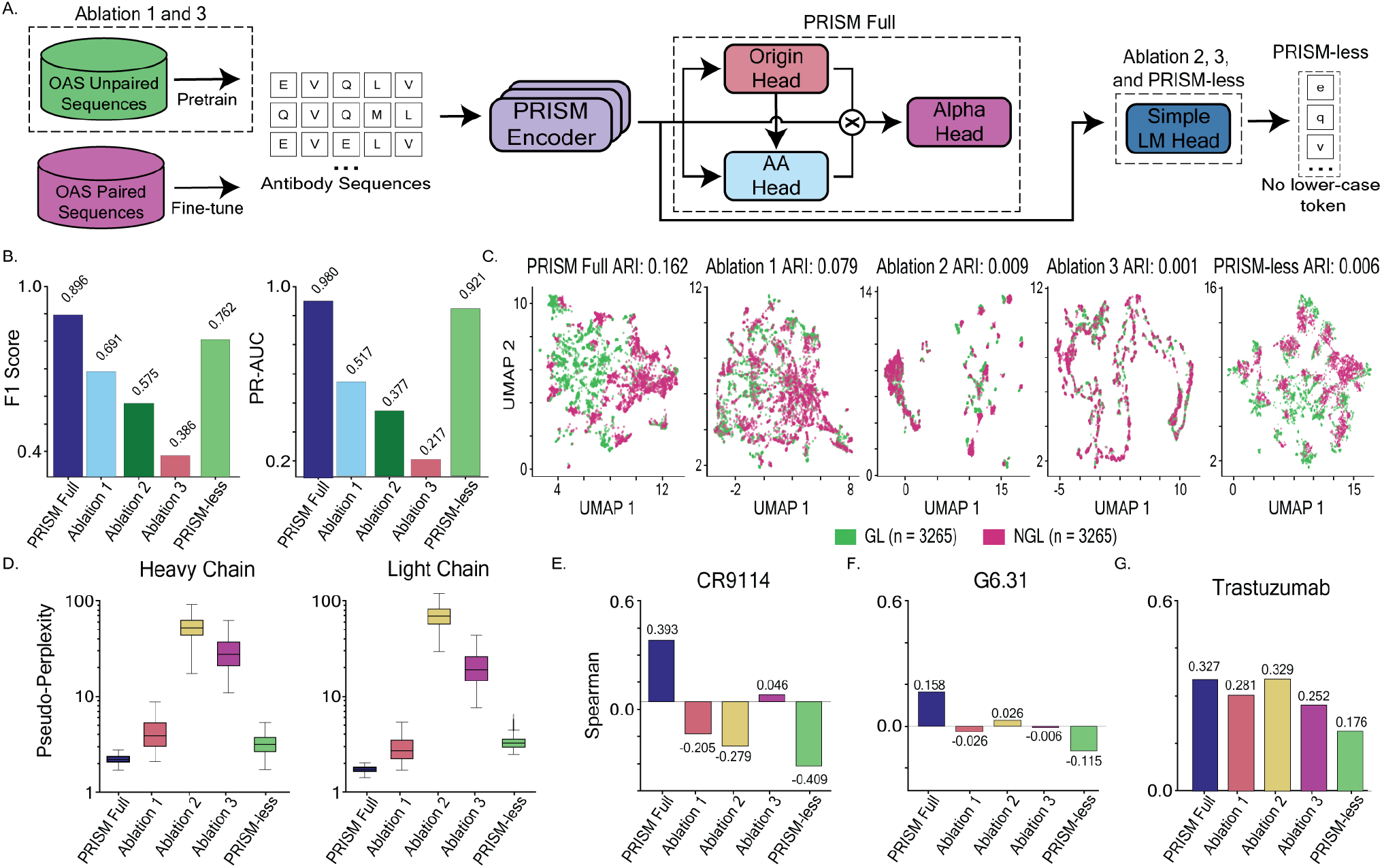
Ablation analysis. **(A)** Comparison of PRISM Full against four ablated variants: Ablation 1 (multi-head, no unpaired pretraining), Ablation 2 (simple LM head, pretrained), Ablation 3 (simple LM head, no pretraining), and PRISM-less (ESM2 initialization with antibody fine-tuning, simple head, upper-case tokens only). **(B, C)** Factorization is essential for GL/NGL separability in the learned representation. **(D)** Impact on generative perplexity across heavy and light chains. **(E-G)** Effect on zero-shot affinity correlations across CR9114, G6.31, and Trastuzumab. PRISM-less attains competitive disentanglement and perplexity but collapses on affinity prediction, isolating architectural factorization—rather than representation quality alone—as the driver of functional performance.

### 7.1. Representational Disentanglement and Sequence Modeling

Architectural factorization and representation quality are separable axes. **PRISM Full** attains near-perfect GL/NGL separability (PR-AUC 0.980, F1 0.896, ARI 0.162), while **Ablation 2** collapses to near-random separability (0.377, ARI 0.009) despite identical pretraining data, confirming that the multi-head is the primary driver of disentanglement; Ablation 1 retains partial separability (0.517/0.079) and Ablation 3 falls to the floor (0.217). **PRISM-less** attains strong probe separability (PR-AUC 0.921, F1 0.762) but near-zero ARI (0.006), indicating ESM2 already encodes a recoverable GL/NGL direction without geometric organization in the embedding space. On generative PPL (Fig. 6D), PRISM Full leads on both chains followed closely by PRISM-less; Ablations 2 and 3 collapse (∼50 and ∼25 heavy-chain PPL) because the simple head cannot map entangled features to the extended GL/NGL vocabulary, while PRISM-less sidesteps this on the standard vocabulary at the cost of forfeiting origin-aware generation (Appendix I).

### 7.2. Zero-Shot Prediction Impact

The functional consequences of factorization are starkest on zero-shot affinity. On CR9114, PRISM Full reaches *ρ*= +0.393 while every ablation fails (Ablation 1: −0.205, Ablation 2: −0.279), and **PRISM-less** produces the strongest *negative* correlation of all variants (−0.409); the same ordering holds on G6.31 (PRISM-less −0.115) and on Trastuzumab (PRISM-less 0.176 vs. PRISM Full’s 0.327). This directly answers whether ESM2 scale plus antibody fine-tuning can substitute for explicit factorization: it cannot. Without an Origin Head to gate predictions, PRISM-less inherits ESM2’s representational competence but defaults to germline bias and systematically penalizes the SHM substitutions that drive binding – representation quality and architectural routing are complementary, not substitutable (Pearson and six developability metrics in Appendix I are strictly consistent).

## 8. Discussion

PRISM mitigates “germline bias” by factoring antibody generation into a joint distribution of evolutionary origin and physicochemical identity, yielding geometrically separable latent representations, controllable generation that independently modulates binding (NGL) and developability (GL), and zero-shot affinity prediction across landscapes where standard ALMs collapse to germline-conservative scoring. Our ablations isolate architectural factorization – not data scale or ESM2 initialization – as the driver of these gains, and we view PRISM as an instance of a broader modeling principle: when sequence generation is governed by a strong hierarchical decomposition (germline scaffold vs. functional diversification, conserved vs. catalytic residues, scaffold-preserving vs. function-modifying mutations), explicitly factorizing the generative process is more effective than relying on a single entangled distribution – particularly in regimes with highly imbalanced signals where rare but functionally important deviations are otherwise suppressed.

### Limitations and Future Directions

Zero-shot developability remains challenging – no model achieves high correlations across all traits from sequence priors alone – and our evaluation is restricted to curated DMS and biophysical panels rather than the full heterogeneous landscape. We plan to fine-tune PRISM on labeled developability data, extend benchmarking to large-scale heterogeneous sets, and leverage the GL/NGL decomposition (treating developability and binding as independent control variables) to resolve the developability-affinity trade-off via reinforcement learning (Blalock et al., 2025).

## Code and Data Availability

Source code, inference scripts, the inference/evaluation dataset, and raw prediction scores for PRISM and all baselines are available at https://github.com/Romerolab/PRISM.

## Impact Statement

This work accelerates therapeutic antibody design. While we acknowledge dual-use risks in generative biology, our focus is strictly clinical and we emphasize the continued importance of ethical guardrails in protein engineering.

## Notes

https://github.com/RomeroLab/PRISM

### Competing Interest Statement

The authors have declared no competing interest.

